# Purifying selection and intraspecies recombination may drive the speciation in Crassostrea: evidence from complete mitochondria sequence of *Crassostrea hongkongensis* and comparative genomic analysis

**DOI:** 10.1101/2024.04.17.589994

**Authors:** Basanta Pravas Sahu, Mohamed Madhar Fazil, Subhasmita Panda, Vengatesen Thiyagarajan

## Abstract

Repeat dynamics and recombination play a crucial role during the evolution of the mitochondrial genome in plants and animals. However, this phenomenon has got less attention within Crassostrea, a complex marine species found worldwide having high commercial value as well as efficient carbon neutralizer. During this study, we characterized the whole mitochondrial genomes of *C. hongkongensis* retrieved from transcriptome data (GenBank acc. no. MZ073671). The current mitochondrial genome (18,616 bp) was composed of a non-coding control region (D-loop region), 2 ribosomal RNA (rRNA genes), 12 protein-coding genes (PCGs), and 23 transfer RNA (tRNA). Furthermore, comparative genomics analysis revealed that the present isolate is closely related to the Chinese isolate (NC_011518) with 99.82% similarity. Microsatellite analysis within the mitochondrial genome revealed its bias towards mononucleotide repeat A/T, di-nucleotide AG followed by AT and AC, trinucleotide AAT followed by AAG, ATC, and ATG. The recombination analysis deciphered the lack of interspecific recombination, but the presence of intraspecific recombination within ND1, ND2, and ND4L of Crassostrea species. Selection pressure analysis revealed the presence of purifying selection within maximum genes which drive the evolution of the species.

## Introduction

Oysters are bivalve mollusks with a wide range of distribution throughout the oceans. Due to their benthic, sessile filter-feeder characteristics, they are considered one of the key drivers for estuary ecology (Bailey & Milner, 2008). Among them, few species are regarded as an excellent resource for aquaculture industries worldwide (Bailey and Milner 2008). Despite their enormous contribution to ecological and economic significance, the information about species diversity and evolutionary history is scanty (Sigwart et al., 2021). Classification of oysters is tedious due to the absence of well-defined morphological characteristics. Species identification through shell morphology remains challenging due to its plasticity to environmental variation (Ghiselli et al., 2021). Few studies suggest Asia Pacific is the epicenter of oyster speciation, but the absence efficient approach to identify them hindered understanding of the evolution of respective species (Ren et al., 2016). Usually, five Crassostrea species, named *C. gigas, C. angulata*, *C. sikamea*, *C. hongkongensis* and *C. ariakensis* are commonly found in China (Liu et al., 2022). In comparison with its related species, *C. hongkongensis* is considered a prized species due to its excellent chevon quality, high market value, and bigger size. Thus, an overwhelming annual production of 1.79 million metric tons was observed along the coast of the South China Sea (Peng et al., 2020). This species is well distributed from the Fujian to Guangxi provinces, along with the densely populated Guangdong province (Ma et al., 2022). It has a great demand in Southern China and contributes to the largest aquaculture production in this area.

Variations in mitochondrial and nuclear genes have been used for phylogenetic analysis. Mitochondrial DNA (mtDNA) is regarded as an excellent molecular marker due to its polymorphism, and maternal inheritance, and thus used to decipher the origin, molecular evolution, and discerning phylogeny of animals (Luikart et al., 2001). Most of the Metazoan mtDNA is circular having 12-13 protein-coding genes (PCGs), 2 ribosomal RNAs (rRNA), and 22 transfer RNAs (tRNA), and the size usually ranges from 14 to 17 kb. The non-coding contains the elements that control the initiation of replication and transcription (Tan et al., 2024). Size variation in mtDNA is usually due to the presence of simple sequence repeats (SSRs) followed by their polymorphism throughout the mitogenome, which can be utilized as repeat-mediated recombination and thus influence the diversity (Jiang et al., 2023; Sharbrough et al., 2023). The presence of SSRs within the mitochondrial genome is well illustrated in many animals (Mayer et al., 2010; Nagpure et al., 2015), and plants (Rajendrakumar et al., 2007; Wynn & Christensen, 2019). However, its information within Crassostrea is limited. Furthermore, both inter and intraspecific recombination observed in several animals, that regulate speciation (Balakirev, 2022; Gantenbein et al., 2005; Tsaousis et al., 2005). However, in Crassostrea, no information is still available concerning the recombination process.

The advent of RNA-sequencing (RNA-Seq) technologies provides a cheap, fast, and efficient strategy to explore gene expression (Metzker, 2010). Except for transcription analysis in eukaryotes, RNA-Seq is also used as an excellent resource for investigating the mitochondrial genome in plants (Forni et al., 2019; Stone & Storchova, 2015) and animals (Marques-Neto et al., 2023; Torson et al., 2022). The aim of our study includes (a) Utilization of RNA-Seq raw data to recover a mitogenome of *C. hongkongensis* and decipher its phylogenetic consequences (b) landscape analysis of simple sequence repeats (SSRs) within the genus Crassostrea (c) investigation of inter and intra specific recombination (d) codon usage and selection pressure analysis to decipher the evolution pattern of those mitochondrial genomes.

## 2. Materials and method

### 2.1. Recovery of mt genome from RNA-Seq

For *in-silico* mining and mitogenome recovery, we have used the RNA seq data previously submitted to Gen Bank from our lab. This data was obtained from the SRA accession number <u>SRX8896407</u>, with the bio project <u>PRJNA643001</u> using a nucleotide BLAST (blastn). The raw RNA-Seq reads were trimmed with Trimmomatic (Bolger et al., 2014) followed by the assembly with Trinity using standard parameters (Haas et al., 2013). Furthermore, RNA-Seq reads were mapped to the reference mitochondrial genome sequences (NC_011518.2), by Bowtie 2, through Geneious v8.1.4 using standard settings (Langmead & Salzberg, 2012). Then assembly of contigs was conducted with BlastN using the reference mtDNA sequences as search queries having an expectation value of 61e10 (Altschul et al., 1990). The annotation was obtained by the MITOS server using the invertebrate animal hereditary code (Bernt et al., 2013) (Table-1, Fig. 1). The tRNA scan-SE version 1.21 was used to find out tRNAs (Laslett & Canbäck, 2008) (Fig. 2).

**Figure 1:**
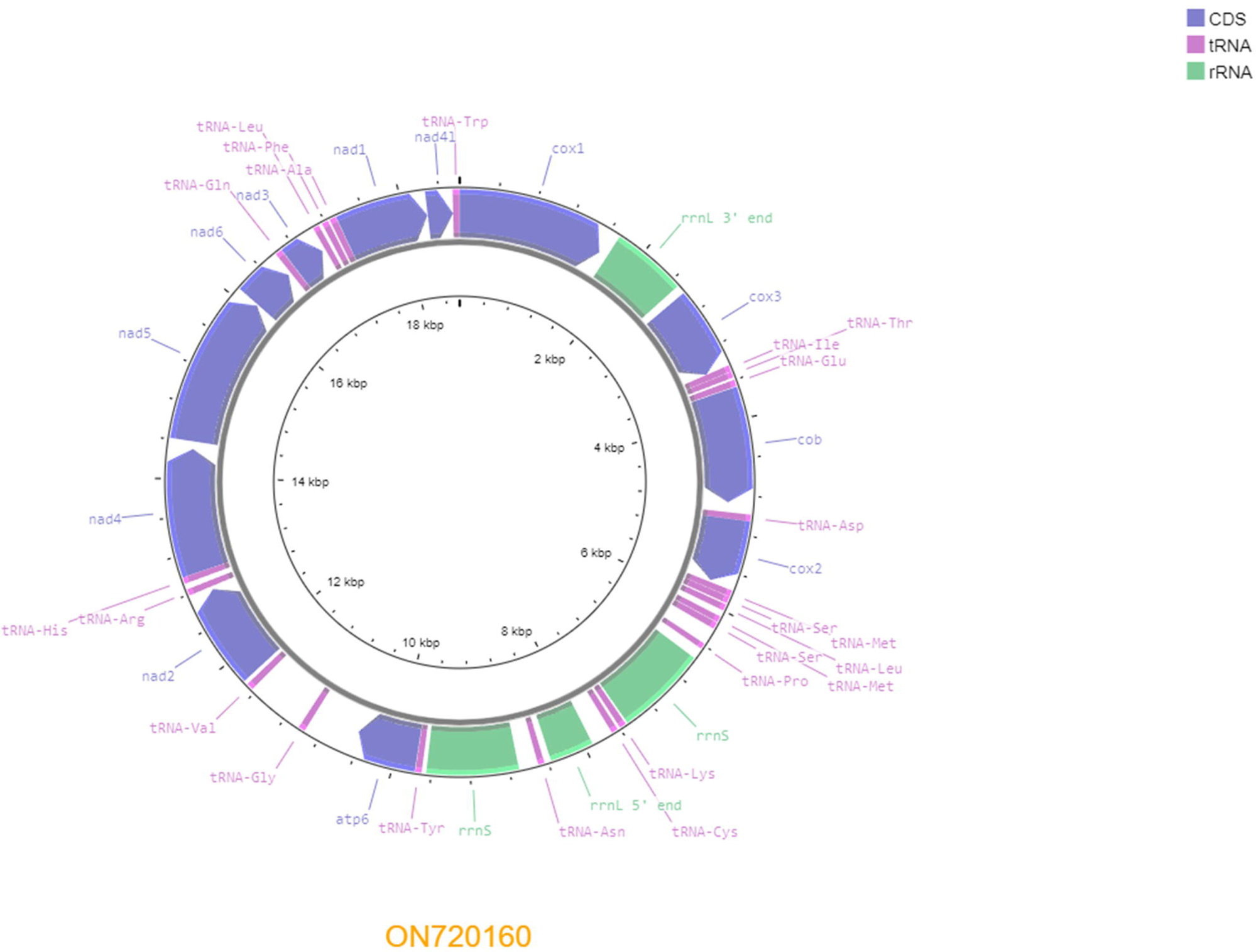
The representation of genetic map of the complete mitogenome of *C. hongkongensis*. tRNA genes are named using single-letter amino acid abbreviations. Inner circle represents the genomic size of the respective genome.

**Figure 2:**
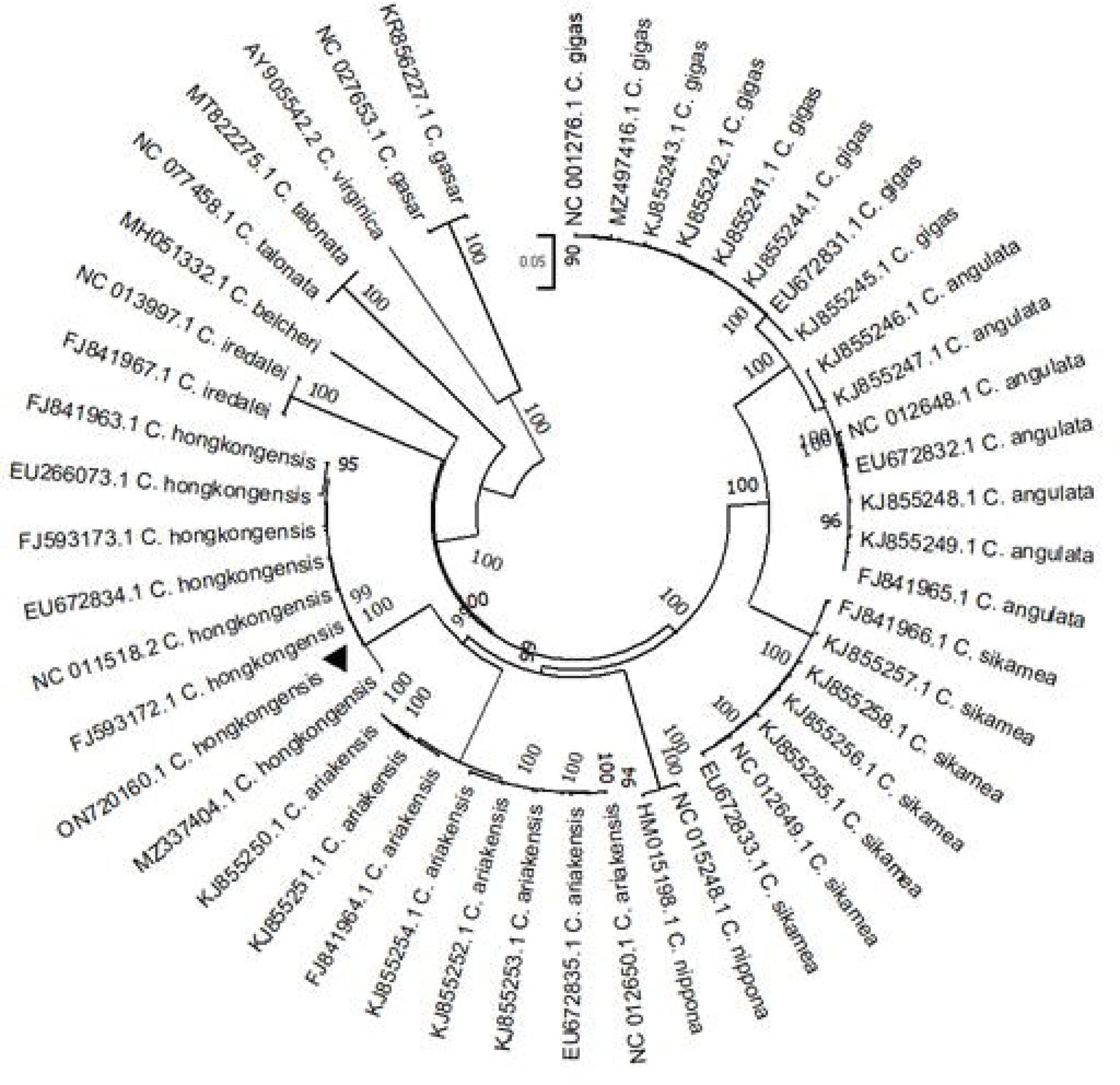
Inferred secondary structure of the 23 tRNAs of the *C. hongkongensis* mitogenome.

### 2.2. Phylogenetic tree and population genetics analysis

To construct the phylogenetic tree, the Neighbor-Joining method of MEGA X along with 1000 bootstrap replications was used (Kumar et al., 2018). The tree was constructed with seven other isolates of *C. hongkongensis* individuals along with forty-one sequences derived from eleven other closely related *C. gigas*, *C. sikamea*, *C. angulata*, *C. ariakensis*, *C. nippona*, *C. belcheri*, *C. gasar*, *C. talonata*, *C. virginica,* and *C. iredalei*. The complete mtDNA was used to obtain the phylogenetic tree (Fig. 3). Population genetics parameters such as parsimony informative sites, number of haplotypes, nucleotide diversity (LJ), haplotype diversity (Hd), selection pressure (dN/dS), and Tajima D calculated by using Dna sp (Rozas et al., 2017).

**Figure 3:**
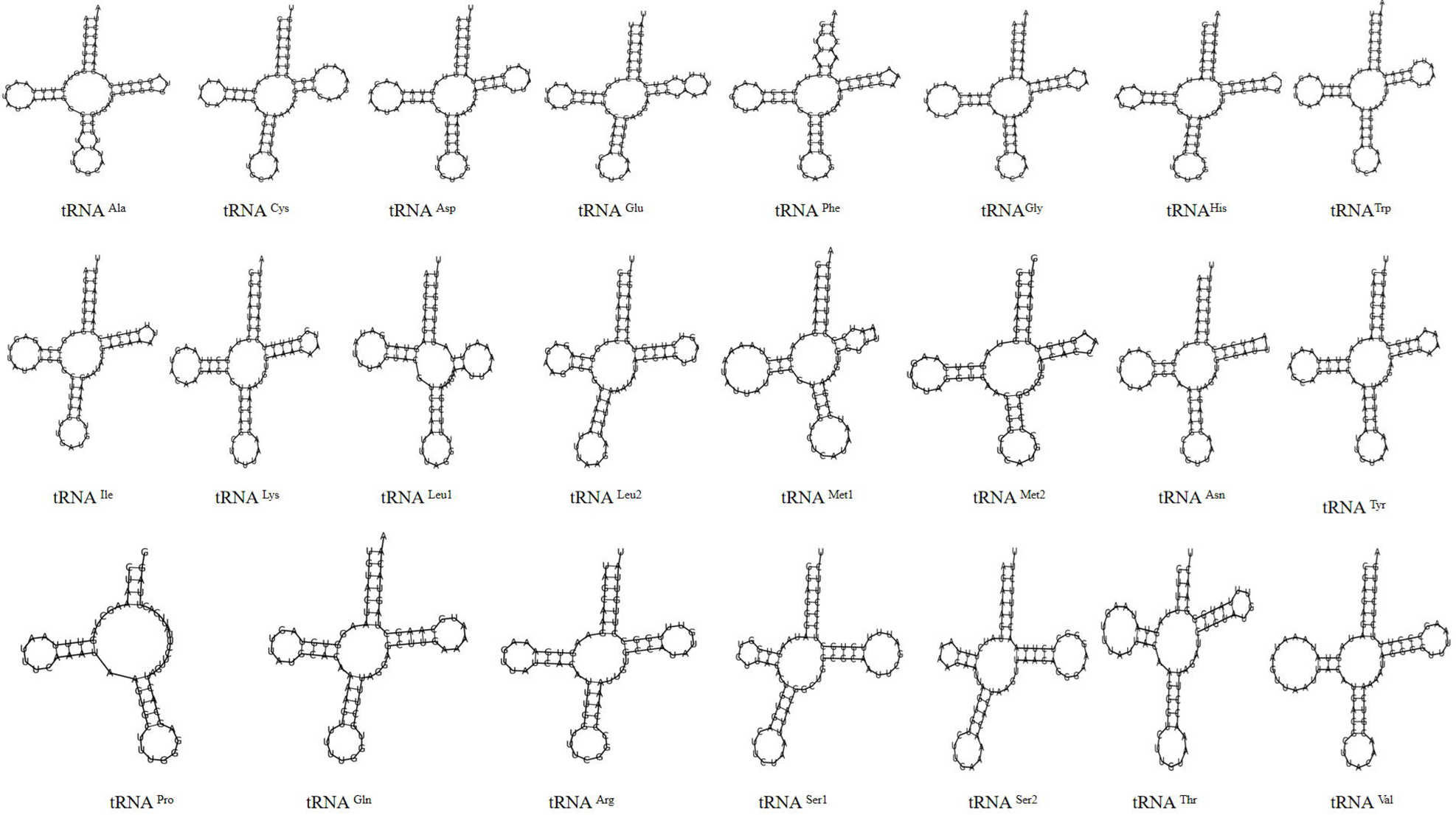
Circular phylogenetic tree based on Neighbor-joining method with bootstrap value 1000 using forty eight complete mitogenome sequences. The accession number along with the species name mentioned at the right side of phylogenetic tree. The ▲ indicates the mitogenome of present study.

### 2.3. SSRs analysis across Crassostrea genus level

The same sequence used for phylogenetic analysis was further used to survey the SSRs. The species along with their accession number were mentioned (Supplementary file-1). The Krait software was utilized to survey the simple and compound SSRs (Du et al., 2018). The parameter for SSR motif searching was set to 6, 3, 3, 3, 3 for mono, di, tri, tetra, and penta, respectively based on previous studies (Nagpure et al., 2015; Sablok et al., 2013). The maximum distance between two consecutive SSRs (dMAX) was 10 nucleotides. Other parameters were set as default.

### 2.4. Recombination analysis

For identification of potential recombinant as well as major and minor parent sequences several recombination detection methods such as RDP, GENECONV, BOOTSCAN, MAXCHI, CHIMAERA, SISCAN, and 3SEQ implemented in RDP4 (Martin et al., 2011). The analysis was carried out with a standard setting with a Bonferroni corrected P-value cut-off of 0.05. RDP datasets include the sequence with 75–90% nucleotide similarity. The position of breakpoints along with the major and minor parents within the genome were manually verified. The recombination events detected by at least three independent methods were only considered (George et al., 2015). Both inter and intraspecific recombination analyses were carried out to identify the major, minor, and recombinant sequences, respectively. Species that have at least three sequences were considered for interspecific recombination analysis.

### 2.5. Codon usage analysis

#### 2.5.1. Nucleotide constraint analysis

The persistence of G and C at the 3rd position of the codon is considered a good indicator of the degree of base composition bias (Zhou & Li, 2009). Thus we checked the reiteration of each nucleotide (A, T, G, C) first (P1), second (P2), and third codon position (P3) within all the PCGs of available Crassostrea mtDNA and revealed their correlation with AT and GC content. The nucleotide bias such as GC skews and AT was measured as 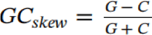 and 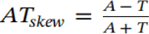

#### 2.5.2. Relative synonymous codon usage (RSCU)

The RSCU is measured by dividing the observed frequency of a codon by the expected frequency if all synonymous codons for that amino acid were used uniformly. The RSCU value of 1 indicates no codon bias for that specific codon, >1.0 indicates positive codon usage bias (defined as abundant codons), and < 1.0 indicates negative codon usage bias (defined as less abundant codons) (Gun et al., 2018; Wang et al., 2018). RSCU value was measured as follows:

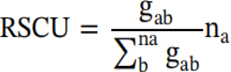

where g_ab_ is the observed number of the *a*th codon for the *b*th amino acid which has na kinds of synonymous codons.

#### 2.5.3. Effective number of codons (ENC)

The ENC value is used to quantify the degree of codon usage bias of a gene irrespective of its length, and its values range from 20 to 61. ENC of a gene < 35 is considered to possess a strong codon bias, whereas > 35 indicates random codon usage (Desingu et al., 2022; Zhou & Li, 2009).

#### 2.5.4. Codon adaptation index (CAI)

The CAI value is used to estimate the level of gene expression (Gupta et al., 2004). Its value ranges from 0 to 1; when the CAI value is close to 1, it indicates a strong bias of the codon usage/protein expression (Tian et al., 2020). The CAI was calculated as follows as follows:

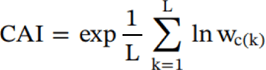

where L is the number of codons in the gene and wc(k) is the ω (relative adaptiveness) value for the k^th^ codon in the gene (Sharp and Li, 1987). To do the above statistical analysis such as nucleotide composition bias, RSCU, ENC, and CAI, the CAIcal server was used (Puigbò et al., 2008).

### 2.6. Analyses of selection pressure on mitochondrial genes

For the identification of positive/negative selected sites within the PCGs of respective mtDNA, nonsynonymous (dN) and synonymous (dS) substitution values of each PCG were calculated through the Datamonkey server with the hyphy package (Weaver et al., 2018). To decipher multiple factors affecting the selective pressure mitochondria gene, the Single likelihood Ancestor Counting (SLAC), followed by the Mixed model of Evolution (MEME) was utilized. The SLAC model can estimate the positive or negative selection, whereas MEME is used to decipher the episodic diversifying selection at each site. For all methods, the cut off p-value was set < 0.05.

## 3. Results and discussion

### 3.1. Characterization of the mitochondrial genome

The complete mtDNA of the current study (ON720160), having a sequence length of 18, 616 bp encodes 12 PCGs, 23 tRNA, two 16S rrnL, and two 12S rrnS (Table-1; Fig. 1). The nucleotide composition revealed its biases towards Adenine and Thymine having nucleotide composition of A: 5398 (29%), T: 4543 (27.30%), C: 6759 (36.31%), and G: 3909 (21%). All the PCGs including tRNA encode by the heavy + strand of mitogenome. COB, NAD3, PCGs begin with ATA as the start codon, while the remaining ten PCGs utilize ATG as a start codon (COX1, COX2, COX3, ATPase6, ND1, ND2, ND4, ND5, ND6, ND4L). Eight PCGs (COX1, COB, COX2, ND1, ND5, ND6, and ND4L) terminated by the TAA codon, while rest four such as ATPase6, ND2, ND3, and ND4 terminated by TAG codon. The 16S rRNA split into rrnL-a and rrnL-b having the length 453 and 772 bp. The 12S rRNA gene was duplicated with a length of 946 bp. The length of tRNA ranged from 62-73 bp, and the tRNACys was the shortest (60 bp) compared with tRNATyr, that composed of the longest nucleotide sequences (73 bp) (Table 1). All the tRNAs observed during our study could be folded into a typical clover-leaf secondary structure except tRNApro, which was lack of one stable DHU (dihydrouridine) arm loop (Fig. 2).

**Table 1:** List of annotated mitochondrial genes in *C. hongkongensis*.

| **Name** | **Start** | **Stop** | **Strand** | **Length** | **Overlapping**<br>**nucleotide** | **Codons**<br>**(Start/Stop)** |
| --- | --- | --- | --- | --- | --- | --- |
| COX1 | 1 | 1617 | + | 1617 | 94 | ATG/TAA |
| 16s rRNA | 1712 | 2483 | + | 772 | 91 |  |
| COX3 | 2575 | 3438 | + | 864 | 0 | ATG/TAA |
| trnI(gat) | 3439 | 3505 | + | 67 | 3 |  |
| trnT(tgt) | 3509 | 3570 | + | 62 | 24 |  |
| trnE(ttc) | 3595 | 3662 | + | 68 | 130 |  |
| COB | 3793 | 4875 | + | 1083 | 108 | ATA/TAA |
| trnD(gtc) | 4984 | 5052 | + | 69 | -14 |  |
| COX2 | 5039 | 5723 | + | 685 | 53 | ATG/T(AA) |
| trnM_0(cat) | 5777 | 5842 | + | 66 | 3 |  |
| trnS1(tct) | 5846 | 5915 | + | 70 | 15 |  |
| trnL2(taa) | 5931 | 5997 | + | 67 | 67 |  |
| trnM_1(cat) | 6065 | 6129 | + | 65 | 6 |  |
| trnS2(tga) | 6136 | 6205 | + | 70 | 1 |  |
| trnP(tgg) | 6383 | 6442 | + | 60 | 117 |  |
| 12s rRNA | 6560 | 7505 | + | 946 | 20 |  |
| trnK(ttt) | 7526 | 7594 | + | 69 | 29 |  |
| trnC(gca) | 7624 | 7691 | + | 68 | 225 |  |
| 16s rRNA | 7917 | 8369 | + | 453 | 71 |  |
| trnN(gtt) | 8441 | 8506 | + | 66 | 198 |  |
| 12s rRNA | 8705 | 9650 | + | 946 | -7 |  |
| trnY(gta) | 9697 | 9762 | + | 66 | 5 |  |
| ATPase6 | 9768 | 10451 | + | 684 | 512 | ATG/TAG |
| trnG(tcc) | 10964 | 11033 | + | 70 | 151 |  |
| trnV(tac) | 11640 | 11712 | + | 73 | 42 |  |
| ND2 | 11755 | 12753 | + | 999 | 34 | ATG/TAG |
| trnR(tcg) | 12788 | 12854 | + | 67 | 60 |  |
| trnH(gtg) | 12915 | 12979 | + | 65 | 44 |  |
| ND4 | 13024 | 14331 | + | 1308 | 83 | ATG/TAG |
| ND5 | 14415 | 16085 | + | 1671 | 11 | ATG/TAA |
| ND6 | 16097 | 16573 | + | 477 | 32 | ATT/TAA |
| trnQ(ttg) | 16606 | 16674 | + | 69 | 2 |  |
| ND3 | 16677 | 17030 | + | 354 | 34 | ATA/TAG |
| trnL1(tag) | 17065 | 17129 | + | 65 | 36 |  |
| trnF(gaa) | 17166 | 17232 | + | 67 | 19 |  |
| trnA(tgc) | 17252 | 17320 | + | 69 | 4 |  |
| ND1 | 17325 | 18260 | + | 936 | 3 | ATG/TAA |
| ND4L | 18264 | 18555 | + | 292 | -12 | ATG/T(AA) |
| trnW(tca) | 18544 | 18612 | + | 69 | 4 |  |

### 3.2. Phylogenetic analysis

The circular phylogenetic tree revealed that the present mitogenome isolate has a close similarity with the MZ337404, that derived from from East China Sea (Fig. 4). However, we observed a species-specific clade within the phylogenetic tree. The genetic diversity analysis using all eight sequences of *C. hongkongensis* mt genome sequences revealed 90 variable sites with 7 haplotypes, Parsimony informative sites: 29, polymorphic sites (S): 90, Nucleotide diversity (LJ): 0.00172, Haplotype (Hd): 0.964, Tajima’s D: −1.00607 (P > 0.10).

### 3.3. SSRs landscape analysis within Crassostrea mitogenome

A total of fourty-eight complete mitochondria sequences from 11 different species of the genus Crassostrea were retrieved from the NCBI database for SSR analysis. All the selected mtDNA with different sizes varying from 17244 - 22446 bp in *C. virginica* and *C. iradale*i, respectively. The overall GC content within the SSRs ranged from 37.13 - 34.65%. The incident frequency of SSRs was lowest in *C. virginica* and highest in *C. iradalei*. A highly variable relative abundance (RA) and the relative density (RD) of SSRs were observed that ranged from 6.742-5.48/kb and 43.79-35.66/kb, respectively. The distribution pattern revealed that 70.29% and 29.71% of microsatellite motifs were distributed within the coding and non-coding regions. From the coding region, the ND1 gene (35.85%) has the highest percentage of SSRs followed by ND2 (30.02%) and ND4l (25.85%). The cSSR analysis revealed a maximum 13 number of cSSRs in *C. iradalei* whereas the lowest *C. angulata* had 4 cSSR in their mtDNA. The cRA and cRD ranged from 0.57 (*C. iradalei*) - 0.21 (*C. angulata*); 8.32 (*C. angulata*) - 3.40/kb (*C. belcheri*), respectively (Figure-). The highest and lowest cSSR% was observed in *C. belcheri* (21.32%) and *C. sikamea* (8.27%). A similar percentage of cSSR distribution was observed within coding and noncoding regions of the respective metagenomes. The incidence of mononucleotide repeat A/T was prevalent compared with G/C thin all analyzed mitochondrial genomes. The dinucleotide repeat AG was highest in *C. belchiri* and lowest in *C. hongkongensis*. Similarly, AC and AT is maximum in *C. belchiri* and *C. iradalei* respectively; whereas the minimum number of AC and AT was found in *C. virginica* and *C. tolonata*, respectively. Overall, repeat motif AG was the most predominant followed by AT and AC. Various types of trinucleotide repeat motifs were scattered throughout the mtDNAs of the Crassostrea species. Among them, AAT was predominant followed by AAG, ATC, and ATG. AAAT tetranucleotide was found only in *C. iradalei* and *C. nipona*. A single pentanucleotide i.e. AAAAC was found in *C. sikamea*. No hexanucleotide was observed in any of the mtDNA of Crassostrea species.

Furthermore, correlation analysis between the genome size and GC content with the incidence, RA, and RD of SSR and cSSR revealed that *C. hongkongensis* had a positive and strong influence on the number of SSRs (P<0.001) and RA (P<0.05), whereas it has no significant correlation with the GC content. In contrast, *C. angulata* has the incidence of cSSR positively correlated with the GC content of the genome (P<0.0001). We also observed that *C. ariakensis* genome size has a positive significant correlation with RA (P=0.02) and RD (P=0.02) of SSR, as well as with GC content of cSSR (P = 0.04).

### 3.4. Recombination analysis

For interspecific recombination analysis, *C. angulata*, *C. areakensis*, *C. hongkongensis*, and *C. sikamia* as they have enough sequence to do the recombination analysis. We did not observe any recombination events during this analysis suggesting a lack of interspecific recombination between species. The intraspecies recombination analysis revealed the presence of three recombination events (Table 2). For this analysis, eleven separate datasets correspond to each gene of the Crassostrea mitogenome. Although each gene experiences similar recombinants, their major and minor parents were inconsistent. For the ND1 gene region, the recombination breakpoints positioned 833-927, where the *C. hongkongesis* acted as the recombinant sequence and *C. gigas* and *C. ariakenasis* acted as a minor and major parent, respectively. The recombination event 2 occurred at the 924-996 bp position of the ND2 region, where *C. hongkongensis* acted as recombinant; *C. sikamea* and *C. gigas* acted as the major parent whereas *C. areakenasis* acted as the minor parent. The final recombination event occurred within ND4L in position 157-243 bp, in which *C. hongkongnsis* was the major parent and *C. gigas* and *C. areakensis* as the minor and major parent, respectively.

**Table 2:**
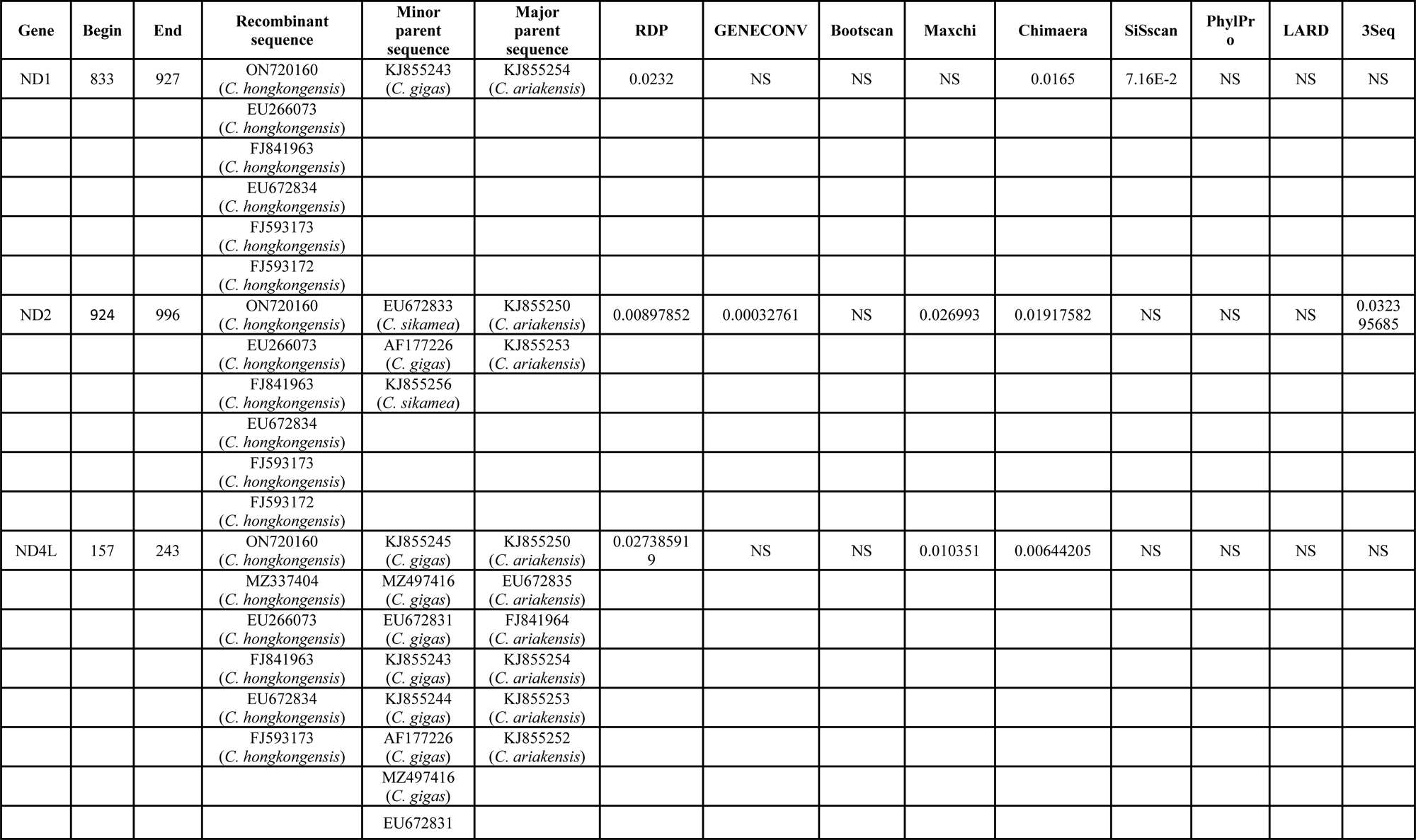
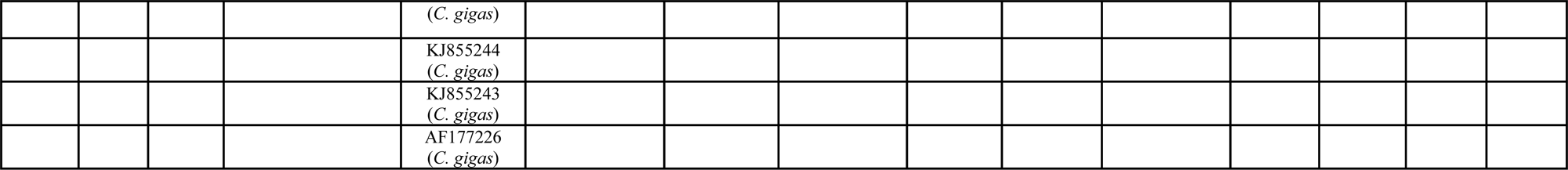
A summary of the recombination events detected within the mitochodrial genome of the genus Crassostrea.

### 3.5. Codon usage analysis

#### 3.5.1. Nucleotide compositional constraint analysis

The nucleotide composition of 12 mitochondrial genes across the species was analyzed. The average value of the nucleotide bases revealed that T (38.47%) was the highest followed by A (24.60%), G (21.96%), and C (14.95%). According to the wobble hypothesis, the 3rd codon position differs considerably in translating 20 different amino acids utilizing the tRNA molecules within the mitochondrial genome. Our analysis revealed the mean value of the nucleotide composition at the 3rd position highest in T3 (42.23%) followed by A3 (28.93%), C3 (18.77%), and G3 (10.04%) among the mitochondrial genes (Supplementary file 2).

#### 3.5.2. Relative synonymous codon usage (RSCU)

The codon usage analysis revealed that all PCGs of Crassostrea mtDNA were enriched with several amino acids such as AAA (Lys=6.33%) followed by ATA (Ile=6.20%), AAT (Asn=5.40), GAA (Asp=4.85), GAT (Ala=5.23), TAT (Asn=4.03), TTA (Leu=4.01), TTT (Phe=3.13), ATT (Ile=3.01). Interestingly all of the major codons were enriched with AT. As all the mtDNA of Crassostrea was enriched with AT, one can expect this pattern of the result within their major codons. RSCU analysis revealed that the majority of codons have both positive and negative codon usage biases. However, the codon-specific RSCU analysis revealed that the positively biased codons were enriched with A/T3 (RSCU>1.0) while negatively biased with G/C3 (Supplementary file 3).

#### 3.5.3. ENC, CAI and their interaction

In the present study, only 2.03% of genes revealed strong codon bias and 97.97% of genes revealed random codon usage. This result was further supported by CAI analysis. The CAI ranged between 0.59-0.80 indicating a moderate gene expression pattern exhibited within mitochondrial genes. So, one can expect some evolutionary forces of mutational bias to create selection pressures that lead to change the diversity to increase fitness in various environments (Supplementary file 4).

### 3.6. Selection pressure analysis

The SLAC analysis revealed that all the codons following purifying/negative selection with the ATPsae 6, COX1, COX2, COX3, ND1, ND2, ND3, ND4, ND5, and ND6. Among them, the COX1 gene exhibited the highest sites (42), followed by ND5 (26) and ND2/ATPase6 (19). The lowest sites were observed within COX2 (6) and ND6 (5) (Table 3). To cross-verify the pattern of selection we furthermore deciphered episodic diversifying selection using all the codons through the MEME method. A total of 35 codons of the twelve genes possessed the episodic selection (Table 4). Among all the genes, a maximum number of episodic selection codons were observed in ND5 (10), followed by COX1/ND2 (4) and ATPase6/COX3/ND1/ND6 (3). Only one codon of ND3 had shown the episodic diversifying selection.

**Table 3:** Selection pressure analysis using SLAC method.

| Gene | Model | Positive/diversifying<br>selection | Negative/purifying<br>selection |
| --- | --- | --- | --- |
| ATP<sub>sae</sub> 6 | Nucleotide GTR(Gamma Time<br>Reversible) | 0 sites | 19 sites |
| COX1 | Nucleotide GTR(Gamma Time<br>Reversible) | 0 sites | 42 sites |
| COX2 | Nucleotide GTR(Gamma Time<br>Reversible) | 0 sites | 6 sites |
| COX3 | Nucleotide GTR(Gamma Time<br>Reversible) | 0 sites | 8 sites |
| ND1 | Nucleotide GTR(Gamma Time<br>Reversible) | 0 sites | 14 sites |
| ND2 | Nucleotide GTR(Gamma Time<br>Reversible) | 0 sites | 19 sites |
| ND3 | Nucleotide GTR(Gamma Time<br>Reversible) | 0 sites | 8 sites |
| ND4 | Nucleotide GTR(Gamma Time<br>Reversible) | 0 sites | 10 sites |
| ND5 | Nucleotide GTR(Gamma Time<br>Reversible) | 0 sites | 26 sites |
| ND6 | Nucleotide GTR(Gamma Time<br>Reversible) | 0 sites | 5 sites |

**Table 4:**
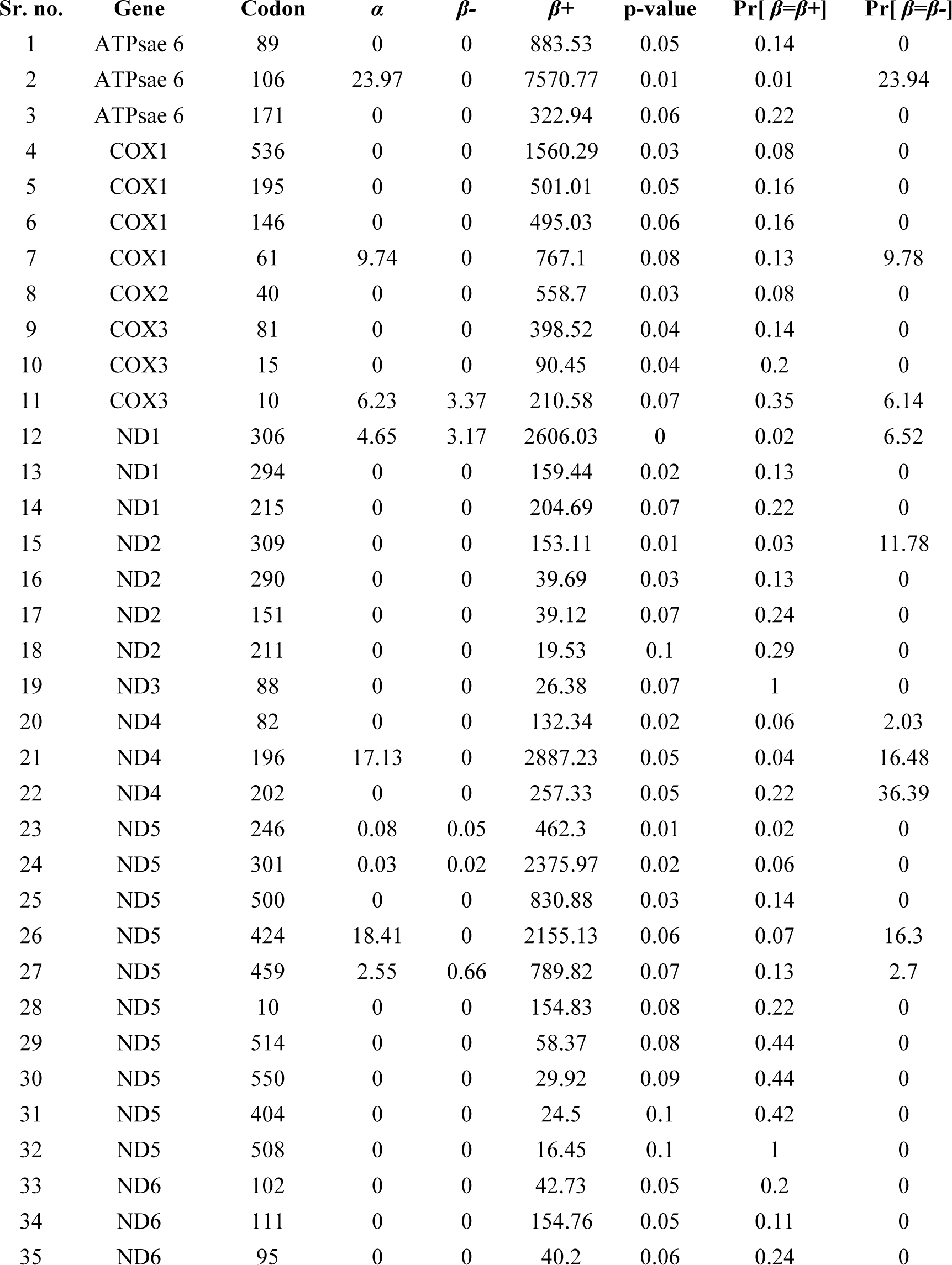
Selection pressure analysis using MEME method.

## 4. Discussion

Recently RNA-Seq data was utilized as an excellent resource to recover various organelle genomes in many organisms (Forni et al., 2019; Torson et al., 2022). A moderate amount of RNA-Seq data could be enough to carry out reference-based assembly or *de-novo* assembly of reads to get the desired mito or chloroplast whole genome. Thus, we employed this strategy to retrieve the *C. hongkongesis* genome from the transcriptome data previously generated from our lab. The reference-based assembly of the mitogenome during the present study deciphered the presence of 12 PCGs, 23 tRNA, and 4 rRNA genes transcribed from a common strand. However, this character is a common feature in marine bivalves along with related species such as *C. gigas* and *C. virginica* (Ren et al., 2016). Although most of the genomic features same as the published mtDNA of *C. hongkongensis*, the genome size as well as the positions of the genes differ. The current mtDNA sequence had a length of 18, 616 bp, which is few base lower than other published sequences. This kind of ambiguity was noticed during the previous study by Liu et al, 2021 in which the deletion of a few bases from intergenic sites reduced the total length of the respective organelle genome and suggested the conserved nature of mitogenome within species (Liu et al., 2022). This phenomenon was further validated by the population genetics parameters and phylogenetic tree. A low level of nucleotide, as well as haplotype diversity along with the clade-specific phylogenetic tree, further strengthens the theory of the conserved nature of mtDNA of the respected species.

Microsatellites are co-dominant molecular markers otherwise known as Simple sequence Repeats (SSRs) or Variable Number of Tandem Repeats (VNTRs) has a wide range of applications such as linkage analysis, population demarcation, parentage assignment, and evolution of the species (Sahoo et al., 2015; Sahoo et al., 2014; Sahu et al., 2014). However, its presence is well documented in the mitochondria of several organisms (Mayer et al., 2010; Nagpure et al., 2015). Within mitochondria, these microsatellites act as a hot spot for mutation, thus leading to the evolution of the species (Jaramillo-Correa & Bousquet, 2005). The genus Crassostrea has got less attention about the presence of microsatellites within its mitogenome along with its type and distribution. Comparative *in-sillico* analysis revealed that the mononucleotide repeats A/T were dominant over the G/C. This type of mononucleotide distribution has been observed in human mitochondria (Chakraborty et al., 2022). However, the chromosome-wide distribution of mononucleotide SSR Crassostrea was exactly the opposite (Sahu et al., 2023). The di and tri-nucleotide SSRs, such as AG and AAT were well conserved within the nuclear as as well as mitochondria of Crassostrea.

Recombination within mitochondria is a natural phenomenon and has been observed within animals, plants, fungi, and protists (Ladoukakis & Zouros, 2017; Rokas et al., 2003). This recombination also leads to hybridization followed by the emergence of sub-species within the lineage (Ladoukakis & Zouros, 2017). Recently, several recombination events were observed within the mtDNA of salangid fishes that led to inter-specific hybridization (Balakirev, 2022). The recombination generally occurs through homologous recombination (intermolecular) or non-homologous recombination (intramolecular) during the recent stages of spermatogenesis (Kraytsberg et al., 2004). Furthermore, repeats are also considered a hot spot for recombination within the mitochondria of plants, animals, and yeast (Balakirev, 2022; Gantenbein et al., 2005; Tsaousis et al., 2005). Therefore, we searched for the presence of recombination within the mtDNA of Crassostrea. Interestingly we did not observe any inter-species recombination events. However, several intraspecies recombination events were observed during this analysis. This type of phenomenon was also noticed during the study of salangid fishes (Balakirev, 2022). Only four out of seven species of Crassostrea actively engaged during recombination. The analysis revealed that all the recombinants were *C. hongkongensis*, while major parents and minor parents were *C. ariakensis* and *C. gigas* in most of the events. We can speculate the hybridization process is going on in nature for the generation of new species. However, it needs further in-depth analysis to prove this hypothesis. Furthermore, only three genes were prone to recombination in our study (ND1, ND2, and ND4l). Intriguingly, during this study, we found these same genes were enriched with repeats as well as recombination events. Does Crassostrea carry out repeat mediated recombination? This argument needs more stringent evaluation in our future research.

Codon usage bias is an evolutionary phenomenon exhibited within all living entities that maintain the selection of the species (Perrière & Thioulouse, 2002). As the mtDNA is prone to a high mutation rate, thus, selection pressure impacts codon composition and vice versa. The codon usage analysis revealed that all PCGs of Crassostrea mtDNA were enriched with many amino acids such as AAA (Lys=6.33%) followed by ATA (Ile=6.20%), AAT (Asn=5.40), GAA (Asp=4.85), GAT (Ala=5.23), TAT (Asn=4.03), TTA (Leu=4.01), TTT (Phe=3.13), ATT (Ile=3.01). Here, most of the codons ended with A/T. Similar codon frequency patterns have been observed in most avian species (Sun et al., 2020). RSCU analysis revealed that codons having positive biases (RSCU>1.0) were enriched with A/T3, while negatively biased genes had G/C3 (RSCU<1.0). However, this type of ambiguity has been observed in Testudines: whose third position is biased towards A/T while in Crocodylia, the mitogenome codons were biased to G/C (Zhou et al., 2016). These examples illustrated the compositional constraints under mutation pressure might have affected the codon usage bias of mitochondrial genes Zhou et al. (2016) (Zhou et al., 2016). A lower ENC value suggests higher gene expression. The ENC value of our study revealed a few genes such as COB, and ND6 that have a great potential for expression. However, a lower to higher ENC value was observed in different mitochondrial genes in vertebrate species such as fish, amphibians, birds, and mammals due to selection pressure (Barbhuiya et al., 2021a, 2021b). Furthermore, the CAI value suggested moderate expression within the Crassostrea. We observed the highest CAI value of 0.79 in the COB gene, followed by a peak of 0.75 in the ND6 gene. It has previously been found that COB adapted towards the use of more efficient codons in the translation of human mitochondria (Levin et al., 2013), as in the case of Crassostrea. The higher translational efficiency of mitochondrial genes in Crassostrea might be related to multiple metabolic factors in response to various environmental conditions.

To decipher the type of selection pressure that drives the evolution of Crassostrea mt DNA, the SLAC followed by the MEME method was utilized. Both methods suggested the presence of negative/purifying selection within the mtDNA. However, this selection was not uniform throughout all the genes. Most of the mutations were found in the COX1 gene followed by followed by ND5 and ND2/ATPase6. Negative selection otherwise known as stabilizing selection by which the organism eliminates the deleterious mutants that lead to adaptation to an environment. Thus, we can expect the prominent mutation within the above-said genes might be responsible for maintaining heterogeneity and increasing the fitness to adapt in extreme environments.

## 5. Conclusion

The genus Crassostrea is a complex species abundantly present throughout the sea with a wide range of environments. The formation of the species, its origin, and its evolution are very challenging. As mtDNA has been evolving faster than nuclear DNA, it can be utilized as an excellent model to disseminate the characteristics of species evolution. Understanding the molecular evolution pattern that was linked with recombination within the current study suggested the presence of hybridization within the species. Further, the purifying selection exhibited within the species of Crassostrea maintains its fitness for diverse environmental situations. However, the presence of several repeats and its association with recombination need further in-depth validation.

## Supporting information

Microsatellite marker characteristics analysis within mtDNA of various species of genus Crassostrea.

Nucleotide constrain analysis during codon usage estimation.

RSCU analysis using all coding region of the genus Crassostrea.

ENC and CAI estimation using all PCGs of the genus Crassostrea

## Supplementary file legends

**Supplementary file 1:** Microsatellite marker characteristics analysis within mtDNA of various species of genus Crassostrea.

**Supplementary file 2:** Nucleotide constrain analysis during codon usage estimation.

**Supplementary file 3:** RSCU analysis using all coding region of the genus Crassostrea.

**Supplementary file 4:** ENC and CAI estimation using all PCGs of the genus Crassostrea.

